# Evidence for an extreme founding effect in a highly successful invasive species

**DOI:** 10.1101/2020.08.04.236695

**Authors:** Kateryna V. Kratzer, Annemarie van der Marel, Colin Garroway, Marta López-Darias, Stephen D. Petersen, Jane M. Waterman

**Affiliations:** Department of Biological Sciences, University of Manitoba, Winnipeg, Manitoba, Canada; Instituto de Productos Naturales y Agrobiología, IPNA-CSIC, San Cristóbal de La Laguna, Tenerife, Spain; Conservation and Research Department, Assiniboine Park Zoo, Winnipeg, Manitoba, Canada

**Keywords:** invasive species, inbreeding, effective population size, genetic bottleneck

## Abstract

The adaptive potential of invasive species is thought to decrease during founding events due to reduced genetic diversity, limiting the new population’s ability to colonize novel habitats. Barbary ground squirrels (*Atlantoxerus getulus*) were purportedly introduced as a single breeding pair to the island of Fuerteventura but have expanded to over a million individuals spread across the island in just over 50 years. We estimated the number of founders and measured the level of genetic diversity in this population using the mitochondrial displacement loop and microsatellite markers. Island samples (*n* = 19) showed no variation in the d-loop, suggesting a single founding female, while Moroccan samples (*n* = 6) each had unique mitochondrial haplotypes. The microsatellite data of the island population (*n* = 256 individuals) revealed a small effective population size, low levels of heterozygosity, and high levels of inbreeding, supporting a founding population size of two to three individuals. Our results suggest that *A. getulus* has undergone an intense genetic bottleneck during their colonization of the island. They are one of the few species where introduction effort does not explain invasion success, although further investigation may explain how they have avoided the worst expected effects following an extreme genetic bottleneck.

## 1. Introduction

Extreme population bottlenecks can produce inbreeding and subsequent inbreeding depression [1, 2] because genetic drift becomes more powerful than selection in small populations. When drift is strong, beneficial alleles can be lost and detrimental alleles fixed due to random chance. As homozygosity increases due to drift, phenotypes associated with deleterious alleles that are hidden in heterozygote states become exposed to selection, and inbreeding depression occurs [for reviews, see 3, 4]. The strength of drift is often not apparent from the census size of a population, as not all individuals contribute equally to the next generation and population size can recover from a bottleneck much faster than the population’s genetic diversity. However, a population experiences drift at the rate of its effective population size, which underscores the fact that even large populations can continue to experience strong effects of drift and continued loss of genetic diversity [5,6].

In some cases, the effects of inbreeding following extreme bottlenecks are not noticeable; thus, understanding the nature of such populations is important for conservation. Within invasive species ecology, many populations are paradoxically founded by a small number of individuals with reduced genetic diversity due to the small size of the available gene pool [7-9]. The ability of these species to adapt to and colonize novel environments can be jeopardized by low levels of genetic diversity [10]. But a sufficiently large founder population (number of individuals or genotypes) [e.g. 11, 12], or multiple introduction events, which introduce new alleles into the population [e.g. 13, 14; see also 7, 15] often characterize successful invasions. Bottlenecked populations that retain sufficient levels of variation may regain some genetic variability through mutation [10, 16, 17], increasing their likelihood of survival. Small founder populations without subsequent introductions should, therefore, have decreased fitness and face difficulty when attempting to establish in novel environments. No successful establishment of an invasive mammal from either one breeding pair or one pregnant female has been recorded; an invasive population founded by either scenario would be an ideal study model for the founder effect [18].

Here we quantify genetic diversity and estimate the effective population size of the invasive population of Barbary ground squirrels (*Atlantoxerus getulus*) on the island of Fuerteventura, Spain. Purportedly introduced as a breeding pair from Sidi Ifni, Morocco in 1965 [19], the current island population has had remarkable success in population growth (estimated one million) and range expansion [20, 21]. We examined the mitochondrial and nuclear diversity of *A. getulus* to resolve any discrepancies between the two differently inherited genomes [22-24]. We targeted the mitochondrial displacement loop and nuclear microsatellites, as any variation in this recently established population would likely be found in the most rapidly evolving areas of the two genomes [25]. We expected to find a single mitochondrial haplotype, high levels of inbreeding, one to four microsatellite alleles at each nuclear locus, and a small effective population size on the island due to the exclusive presence of alleles from a single founding pair. With this research, we intend to contribute to the general knowledge on the role of genetic diversity and bottlenecks in explaining the success of biological populations.

## 2. Methods

### (a) Study species, trapping locations and methods

We trapped *A. getulus* according to previously described methods in various locations on Fuerteventura and Morocco [see 26-29] and stored tissue samples in 95% ethanol. Mitochondrial d-loop sequences were obtained from 45 animals, and 256 animals were genotyped at eleven microsatellite loci (see S.I. for details).

We tested for inbreeding and variation from Hardy-Weinberg equilibrium using the “adegenet” package v.2.1.1 [30, 31] and the “pegas” package x.0.11 Monte Carlo exact test with 1000 replicates [32], respectively, in R v.3.5.1 [33]. Alleles were determined to have been introduced by founders rather than mutation (i.e. “founding alleles”) if they had a frequency > 0.05 and were more than one repeat unit away from a common allele [12]. We performed a principal component analysis (PCA) using the “ade4” package v.1.7-13 [34] to determine whether there was any genetic structure in the population. We then calculated effective population size (N_e_) using the LDNE method, assuming random mating and setting the minor allele frequency to 0.05 [35].

## 3. Results

### (a) Mitochondrial DNA

We found no variation among island squirrels, whereas all six individuals from Morocco had unique haplotypes and showed 16 variable nucleotide sites compared to island samples, despite the limited sample size of the Moroccan source (Fig. 1, Table 1). We found four variable sites (0.389%) between Fuerteventura sequences and M10, the Moroccan sequence most similar to those on the island (Fig. 1).

**Table 1.** Nucleotide differences within the mitochondrial d-loop of six *Atlantoxerus getulus* haplotypes from Sidi Ifni, Morocco.

| Position | 27 | 95 | 98 | 111 | 114 | 173 | 252 | 258 | 276 | 282 | 289 | 299 | 316 | 746 | 824 | 970 |
| --- | --- | --- | --- | --- | --- | --- | --- | --- | --- | --- | --- | --- | --- | --- | --- | --- |
| M03 | C | A | C | T | G | C | C | C | C | C | A | A | T | A | T | T |
| M06 | . | . | . | . | . | . | T | . | . | . | . | . | . | G | . | . |
| M10 | . | G | . | . | . | . | . | . | T | . | G | G | . | . | C | . |
| M18 | T | . | . | C | . | . | . | . | . | . | . | . | C | . | C | C |
| M19 | . | . | A | . | A | T | . | . | . | . | . | . | . | . | C | . |
| M21 | . | G | . | . | . | . | . | T | . | . | . | . | C | . | C | . |
| FV | . | . | . | . | . | . | . | . | . | T | . | G | . | . | C | . |

**Figure 1.**
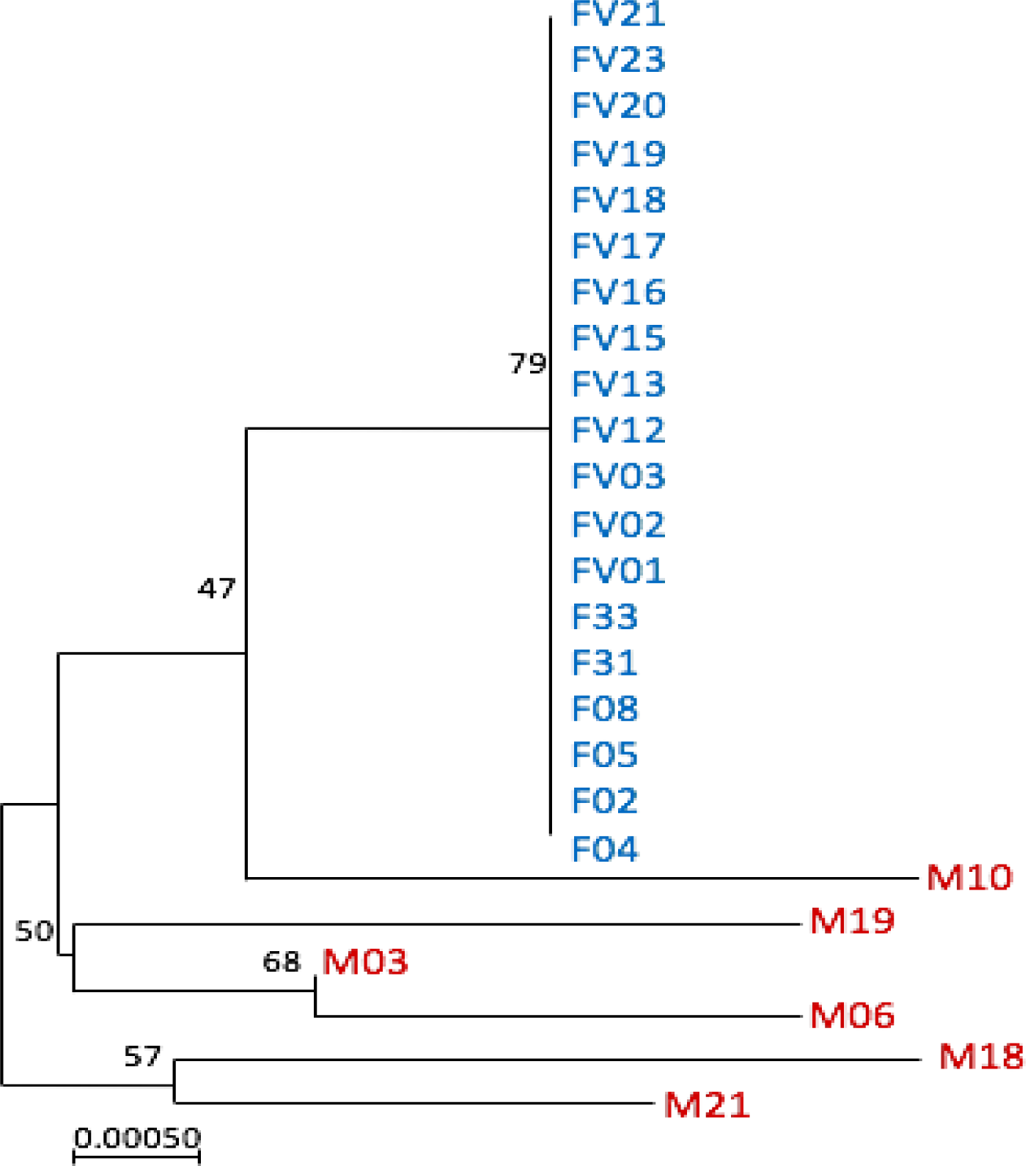
Evolutionary relationships between island (blue) and mainland (red) *Atlantoxerus getulus* based on the mitochondrial DNA displacement loop. Relationships inferred using the Neighbour-Joining method with 1000 bootstrap replicates. Evolutionary distances calculated using the Tamura-Nei method. All codon positions were included (total 1027 positions). Made in MEGA7 [53].

### (b) Nuclear DNA

We found no evidence of large allele dropout or scoring error due to stuttering [36]. Null alleles, indicated by homozygote excess, were present at five loci that were removed from the analysis [37, 38]. All remaining loci were in HWE (*p* > 0.05). Each locus had between two and nine alleles (4.36 ± 2.11, mean ± SD), the number of founding alleles ranging from one to five (2.73 ± 0.65). Mean observed heterozygosity (H_O_ = 0.57) was greater than expected (H_E_ = 0.55; Table 1 supplemental information) and the average level of inbreeding was high (Fig. 2: average *F* = 0.23 [0.10 – 0.60, min - max]). Since we found no evidence of population structure (S.I. Fig. 1), we assumed that our sample was representative of the entire island population. We estimated N_e_ to be 77.2 (95% CI: 56.3, 109.5).

**Figure 2.**
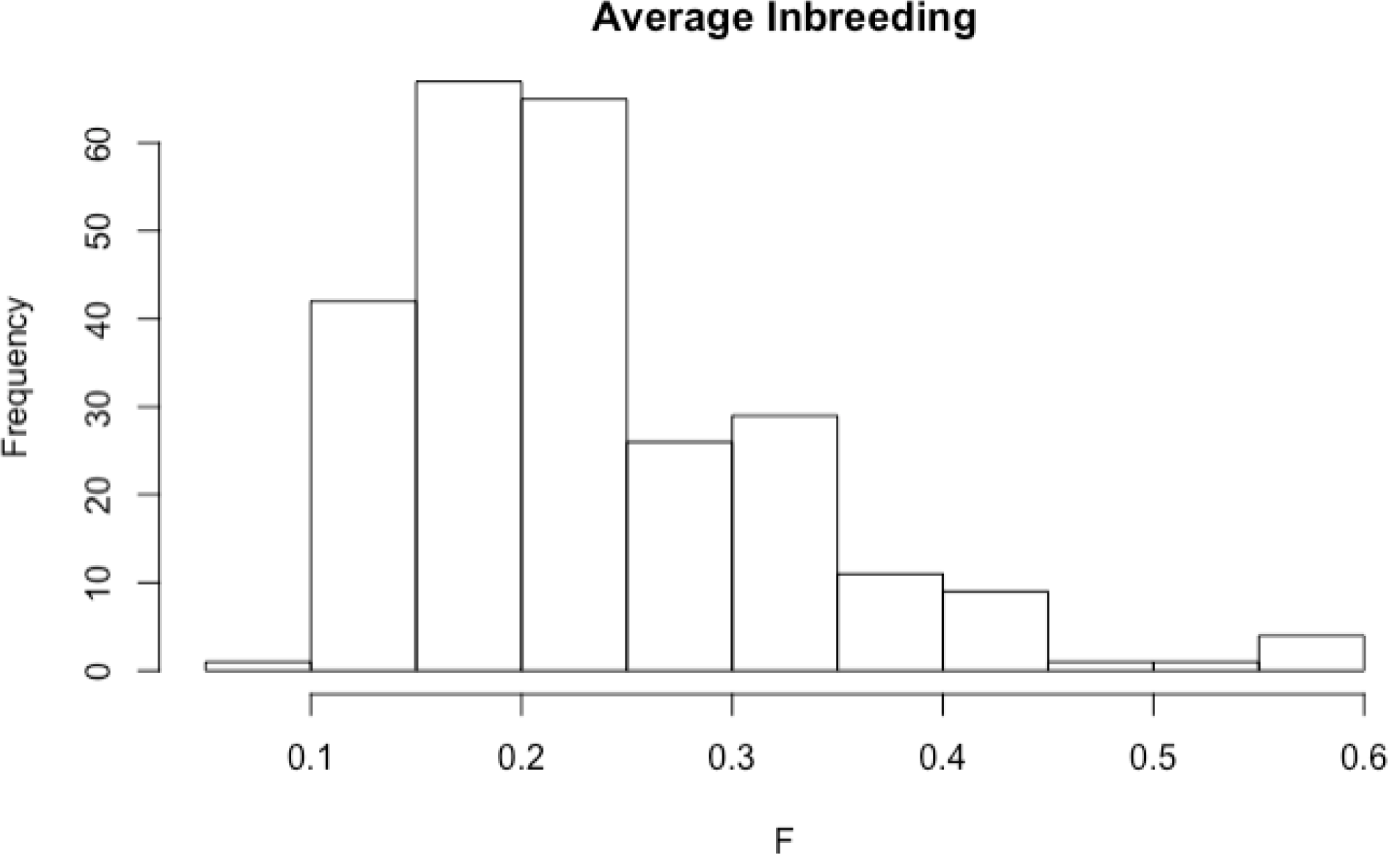
Average inbreeding coefficients of 256 *Atlantoxerus getulus* individuals based on microsatellite markers of nuclear DNA. The *F* values ranged from 0.097 – 0.596 (mean *F* = 0.233).

## 4. Discussion

We characterized segments of the mitochondrial and nuclear genomes of a highly successful invasive island population of *A. getulus* to determine its genetic diversity and number of founders. We observed low genetic diversity, evidence of inbreeding in mitochondrial and nuclear DNA, and a single mitochondrial haplotype suggesting the presence of only one founding female. We found variation between each mitochondrial d-loop sequence of Moroccan samples despite a small sample size (*n* = 6), whereas the island population did not show variation with a larger sample size (*n* = 19).

Microsatellite data also supported the hypothesis that this island population was founded by a small number of individuals but data from marker Aget19 suggest that there may be more than two founders (S. I. Table 2). Of nine alleles at this locus, five are present at a frequency greater than 5% [12], which is incongruous with the hypothesis that the island population was founded by two individuals. However, two of these alleles (repeat lengths 319 and 339) have frequencies just above the threshold of being counted as true founder alleles (0.0573 and 0.0553, respectively; S. I. Table 2). It is possible these alleles were introduced by a founder, but the potential that 319 and 339 are due to rare double mutations, genotyping error, or an early mutation that was propagated over the threshold cannot be overlooked. Another microsatellite marker, Aget1, also has a high number of alleles but only two are present at high frequency (> 5%). An interesting allele at this marker is repeat length 152, which is two repeat units away from a founder allele and therefore does not comply with the recommended criteria [12]. However, it is present at low frequency (0.0108), and multiple mutations in the same location, while unlikely, are not impossible [12]. Further investigation may confirm the true origin of these alleles.

With an average inbreeding coefficient *F* of 0.23, the *A. getulus* population should be at a survival or range expansion disadvantage [39, 40], as an increased probability of extinction exists when *F* values are at or just below “intermediate” levels (0.30 – 0.40; 11, 41]. However, the species has successfully established and spread across the island [20, 21] in a genetic paradox of invasion [9]. *Atlantoxerus getulus* invasion success may be due to extrinsic habitat factors [22, 29], or other species-level [42, 43], behaviour [28], or life-history traits [44]. Alternatively, inbreeding may have benefitted the population by purging deleterious founding alleles [41, 45]. Despite an estimated population size of one million, the effective population size was approximately 77 individuals (0.0077%), which is very low compared to other infamously bottlenecked mammals. Northern elephant seals survived near extinction and experienced steady population growth from about 100 to over 200,000 individuals, with an N_e_ of approximately 40,000 (>20%) [12, 46, 47]. Cheetahs are estimated to number 6674 individuals with an N_e_ of between 1001–2937 (15–44%) [48, 49]. Some re-introduced populations of European bison (*Bison bonasus*) have N_e_/N values as low as 0.05 (5%) [50]. The island population of *A. getulus*, therefore, has one of the smallest recorded effective population sizes relative to their census size.

One caveat of our study was the sampling regime. The sampling density for mtDNA was low although samples were collected from sites across the entire island of Fuerteventura, whereas sampling density for nuclear DNA was higher but restricted to a single area. As such, we found no evidence of population structure. However, there are no geographic barriers to dispersal across the island, as squirrels have been observed in all regions [20, 21], thus population structure may be absent altogether. Better coverage of the island or perhaps the collection of whole genomes may provide further insight into this recent founding event.

We have shown that the *A. getulus* population on Fuerteventura has undergone an intense genetic bottleneck during their colonization of the island. However, despite their lack of genetic diversity and low effective population size, they have successfully established and spread across the island, providing an ideal example of the founder effect.

## Ethics

All sample collection followed the animal care protocols of the University of Manitoba (Animal Care and Use Committee #F14-032) and the government of Fuerteventura (Cabildo Insular de Fuerteventura #14885). Samples from Morocco were obtained with the permission of the Ministry of Territory Development, Water and Environment of Morocco (512/0170 March 2006) and brought back to the EU under the authorization of the Government of the Canary Islands.

## Data accessibility

Data will be available at Dryad

## Authors’ contributions

JMW and ML-D discussed the idea and KVK, JMW and SDP designed the study. KVK, AVDM, ML-D and JMW conducted the field work. KVK and AVDM performed the lab work. KVK, CG, and SDP contributed to the analysis and interpretation of the data. All authors contributed to the manuscript preparation and revisions. All authors approved the final version of the manuscript and agree to be held accountable for the content of this publication.

## Acknowledgements

We would like to acknowledge Lucy Johnson for indispensable assistance and instruction in laboratory and analysis techniques.

