## Supplemental methods figures tables for "Evidence for an extreme founding effect in a highly successful invasive species"

**Supplemental Information**

**Methods**

**(a) Study species**

*Atlantoxerus getulus* are social with females living in matrilineal kin groups, while adult males nest with unrelated males and subadults of either sex [51]. Their life history traits fit on the fast side of the fast-slow life history continuum [44]. Most males and females breed within the first breeding season after birth and breeding can occur annually or biannually [44]. On average 78% of females are successful in raising their litter (average 3 offspring; range 1-8) to emergence, although offspring mortality ranges between 20 and 50 % [44]. The lifespan of the surviving individuals ranges from 1.5 to 5 years [44, 52].

**(b) Genetic analysis – mitochondrial DNA**

We extracted DNA from 85 samples using the E.Z.N.A.® Tissue DNA Kit (Omega Bio-tek, Norcross, GA), from which 45 were successfully amplified and sequenced (34 island, 11 mainland). We included 25 (19 island, six mainland) complete displacement loop sequences in our mtDNA analysis. We used four primers to target the d-loop (S. I. Table 3). We aligned d-loop sequences using the automatic ClustalW parameters in MEGA7 [53]. Samples with missing or ambiguous nucleotide data were removed from the analysis, and d-loop sequences were trimmed to a uniform length using the most closely related reference sequence available (*Spermophilus dauricus*, GenBank NC_027283.1).

**(c) Genetic analysis – nuclear DNA**

We isolated the genomic DNA of 256 individuals using Chelex beads (Bio-Rad) [54].Two microsatellite primers were previously designed for Cape ground squirrels (*Xerus inauris*; Xin09 and Xin10) [55] and we newly developed nine primers, which we labelled with fluorescent dyes and pooled according to annealing temperatures [51]. We examined nuclear DNA using GeneMarker (SoftGenetics, State College, PA, USA), which scored the loci semi-automatically. At least two observers scored the alleles in an objective scoring procedure [56]. We conducted tests for null alleles using Micro-Checker [36].

**Figures and Tables**

**Supplemental Information Table 1.** The characterization of eleven microsatellite loci for Barbary ground squirrels (*Atlantoxerus getulus)*. Annealing temperature of primers (T_A_), number of alleles (N_A_), and observed and expected heterozygosities (H_O_/H_E_) are given.

| Locus | ^Dye label^Primer sequence | Repeat motif | T_A_ (˚C) | N_A_ | Allele  size | H_O_ | H_E_ |
| --- | --- | --- | --- | --- | --- | --- | --- |
| Aget1 | ^PET^AAACCCTATCTTCCTTCTATGAGC | AGAT | 55 | 8 | 160-175 | 0.59 | 0.60 |
| Aget3 | ^FAM^AAATACAAATCCACACACAAGAGG | AAAAG | 55 | 2 | 168-178 | 0.45 | 0.40 |
| Aget4 | ^NED^GCTGTGCTCTCTTCATGATTCTC | ACAG | 55 | 4 | 180-195 | 0.44 | 0.45 |
| Aget8 | ^PET^ACAGGAGCCCATTTGTATATGTC | ATCC | 55 | 4 | 230-240 | 0.70 | 0.68 |
| Aget17 | ^FAM^AGACATGATCACTTAATTCCCTCC | AGAT | 55 | 5 | 320-374 | 0.73 | 0.73 |
| Aget19 | ^VIC^ATTCCCTCCAAGATATCCATCCC | AAAG | 55 | 9 | 310-332 | 0.74 | 0.71 |
| Aget23 | ^FAM^TGGAATGTTTGGCCATATGTGAG | AGAT | 60 | 4 | 370-384 | 0.53 | 0.49 |
| Aget34 | ^FAM^GGGATGCTTTAATACTGAGTCCC | AGAT | 55 | 3 | 473-485 | 0.59 | 0.55 |
| Aget42 | ^FAM^AAACCCTTCTTATCACATGCACC | AGAT | 55 | 5 | 320-375 | 0.67 | 0.68 |
| Xin09^1^ | ^FAM^CCTCATCACAACCAAGACAG | GT | 55 | 4 | 160-170 | 0.31 | 0.29 |
| Xin10 | ^FAM^CAGATTGAGAGTGAGAGGTG | GT | 55 | 6 | 212-235 | 0.49 | 0.50 |

^1^Xin loci were developed for the closely related Cape ground squirrel (*Xerus inauris*) [55]. ^2^The step-up thermocycle profile was 95˚C for 5 minutes (1x), then a cycle of 94°C for 30s, the step-up (45°C 10 times 30s, 55°C 20 times for 30s, and 72°C for 30s (32x)), ending with 72°C for 30 min and an infinite hold at 8°C (1x).

**Supplemental Information Table 2.** Allele frequencies for eleven microsatellite markers in an island population of Barbary ground squirrel (*Atlantoxerus getulus*).

| **Locus** | **Allele (Repeat Length)** | **Allele Frequency** |
| --- | --- | --- |
| Aget1 | 152 | 0.0108 |
|  | 156 | 0.0269 |
|  | **160** | 0.3925 |
|  | 164 | 0.0054 |
|  | 168 | 0.0403 |
|  | **172** | 0.4946 |
|  | 176 | 0.0296 |
| Aget3 | **169** | 0.7198 |
|  | **179** | 0.2802 |
| Aget4 | 184 | 0.0061 |
|  | **186** | 0.1061 |
|  | **192** | 0.7082 |
|  | **194** | 0.1796 |
| Aget8 | 226 | 0.0286 |
|  | **230** | 0.3194 |
|  | **234** | 0.3590 |
|  | **238** | 0.2930 |
| Aget17 | **322** | 0.2183 |
|  | **362** | 0.2664 |
|  | **370** | 0.3515 |
|  | **374** | 0.1638 |
| Aget19 | 311 | 0.0119 |
|  | **315** | 0.3992 |
|  | ***319*** | *0.0573* |
|  | 323 | 0.0059 |
|  | 327 | 0.0059 |
|  | **331** | 0.1146 |
|  | **335** | 0.3360 |
|  | ***339*** | *0.0553* |
|  | 343 | 0.0138 |
| Aget23 | 372 | 0.0109 |
|  | **376** | 0.1095 |
|  | **380** | 0.1971 |
|  | **384** | 0.6752 |
|  | 388 | 0.0073 |
| Aget34 | **474** | 0.5952 |
|  | **482** | 0.1138 |
|  | **486** | 0.2910 |
| Aget42 | **332** | 0.4588 |
|  | **336** | 0.2654 |
|  | 348 | 0.0165 |
|  | **352** | 0.0658 |
|  | **356** | 0.1934 |
| Xin09^1^ | **163** | 0.1726 |
|  | **169** | 0.8274 |
| Xin10 | **218** | 0.4390 |
|  | **232** | 0.5549 |
|  | 240 | 0.0061 |

Alleles likely introduced to the population by migration (“founding alleles”) rather than mutation are indicated in bold. The two alleles of the Aget19 locus that are slightly above the threshold frequency of 0.05, potentially indicating a larger founding size, are italicized.

**Supplemental Information Table 3.** Primers targeting the displacement loop of mitochondrial DNA in *Atlantoxerus getulus*.

| **ID** | **Sequence** | **Source** |
| --- | --- | --- |
| CB3R-F | 5’ – CAT ATC AAA CCA GAG TGA TAT TTC | Developed from [57] |
| CSBint-R | 5’ – CGT GAA ACC AGC AAC CCG CT | [58] |
| F5-F | 5’ – CAT TAA TAA TGA TGA AAG TAC ATA G | Newly developed |
| 12SAR 120bp-R | 5’ – AGC TCG CCC ATC CAT GCA GCT CA | Developed from [57] |

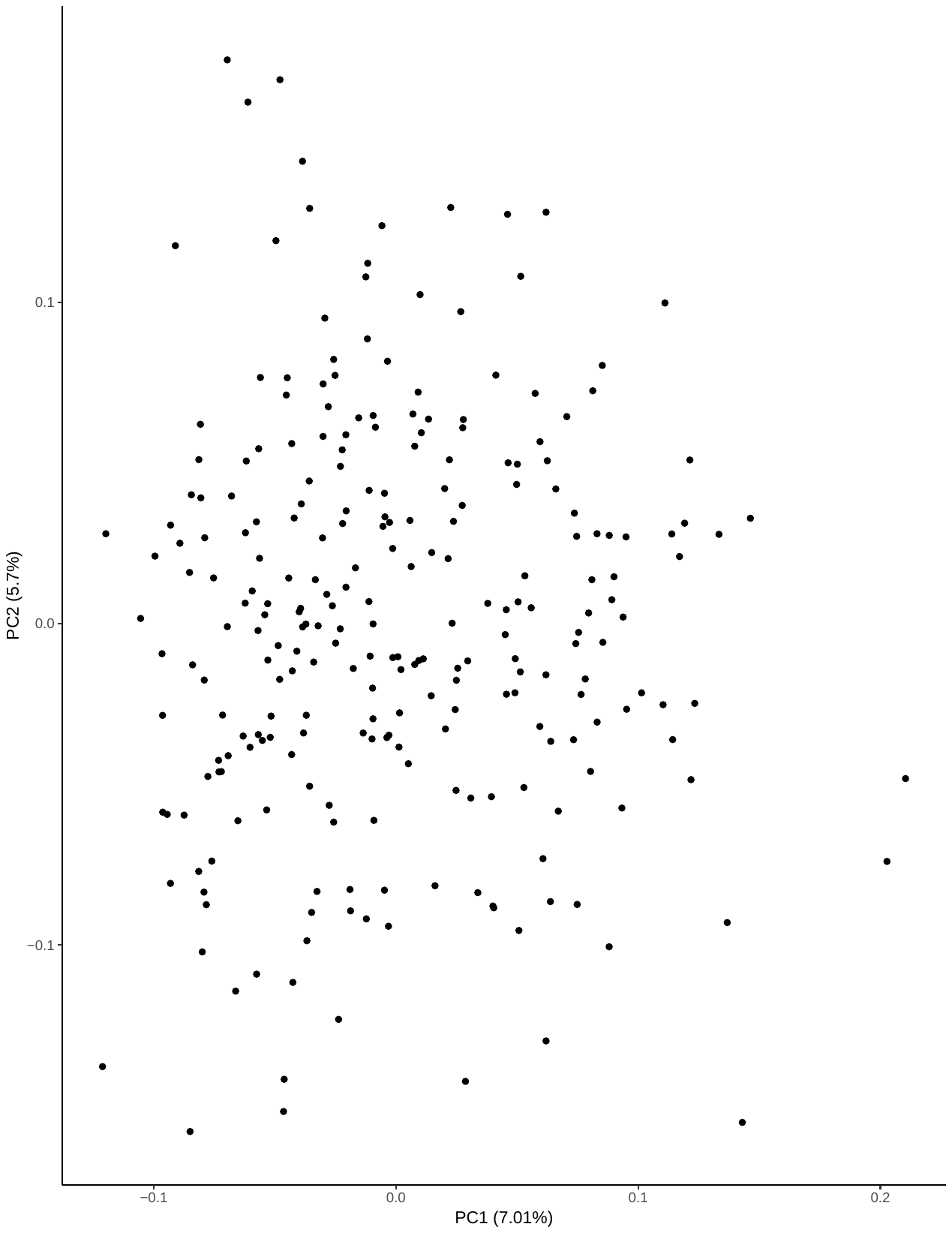

**Supplemental Information Figure 1.** Autosomal population structure of the Fuerteventura population *Atlantoxerus getulus* based on plotting principal components 1 (PC1) and 2 (PC2).
